# Separating robust from fragile cell-type changes in spatial transcriptomics using *screening label uncertainty to identify composition effects*

**DOI:** 10.64898/2026.11.24.768995

**Authors:** Rebecca Sakko, Sakthivel Ravi, Holden E. Harris, Jose F. Abisambra

**Affiliations:** Center for Translational Research in Neurodegenerative Disease (CTRND), University of Florida, Gainesville, FL 32610, USA; Department of Neuroscience, University of Florida, Gainesville, FL 32610, USA; McKnight Brain Institute, University of Florida, Gainesville, FL 32610, USA; Brain Rehabilitation Research Center, Malcom Randall VA Medical Center, North Florida/South Georgia Veterans Health System, Gainesville, FL 32608, USA; Brain Injury Rehabilitation and Neuroresilience (BRAIN) Center, University of Florida, Gainesville, FL 32610, USA; Nature Coast Biological Station, Institute of Food and Agricultural Sciences, University of Florida, Cedar Key, FL 32625, USA

**Keywords:** spatial transcriptomics, Xenium, cell-type annotation, classification confidence, sensitivity analysis, traumatic brain injury

## Abstract

Imaging-based spatial transcriptomics platforms such as 10x Xenium measure thousands of genes in intact tissue at single-cell resolution, making them attractive for screening which cell populations change between experimental conditions. In small studies, however, apparent changes in cell-type composition can arise from misannotated cells rather than biology, and the reliability of individual cell-type calls is rarely reported. We present SLUICE (Screening Label Uncertainty to Identify Composition Effects), an R/Seurat workflow that classifies each cell by atlas-informed marker scoring and records a per-cell confidence margin and runner-up identity. Each group difference is then tested against three diagnostics: region-specific confidence thresholds, sensitivity analyses that reassign borderline calls, and single-gene driver checks that combine gene co-detection, runner-up identity, and spatial location. Applied to Xenium Prime 5K data from six mice (three traumatic brain injury, three sham) in two contrasting regions, the hippocampus and the optic tract, confidence thresholds differed almost sevenfold between regions (0.17 vs 0.026), showing that a fixed cutoff does not transfer between tissues. Driver checks identified mislabelled populations that margin-based filtering left in place, including Klotho (Kl)-expressing neurons called choroid plexus in the hippocampus and Csf3r-expressing myeloid cells called neutrophils in the optic tract. Excluding the neutrophil panel reduced the number of optic tract categories whose group difference changed direction from five to two (one of which was the Unclassified category), while the major glial populations were stable across all analyses. SLUICE separates robust from fragile composition changes, identifying which shifts merit experimental follow-up before time and tissue are committed to them. Code is available at https://github.com/rebeccasakko/hippocampus-xenium-analysis.

## Introduction

Imaging-based spatial transcriptomics now measures hundreds to thousands of genes in intact tissue at single-cell or subcellular resolution, making it an attractive way to screen for cell populations that change between experimental conditions. In injury models such as traumatic brain injury (TBI), where glial and immune populations change in number and state, a spatial screen can show which cell types shift and where they sit relative to the lesion, before more targeted experiments are chosen^1–3^.

The weak point of such a screen is the cell-type label. Each cell must be assigned an identity from a limited gene panel, by clustering followed by manual annotation, by marker scoring, or by mapping to a reference atlas^4,5^. These assignments involve researcher choices that can change the result, such as clustering resolution, marker lists, and score thresholds. Imaging-based data add a further source of ambiguity, because cell segmentation can assign transcripts from neighbouring cells to the wrong cell, which blurs the expression profile that the label depends on^3,6^. Most workflows report one label per cell and give little indication of how close each call was to a different label.

When a few animals per group are compared, a category that holds a few percent of cells can move between conditions because borderline cells were assigned to it. Confidence scores from reference-mapping tools can help, but a single fixed cutoff assumes that the score means the same thing in every tissue. Treating the label as certain, or filtering with one cutoff and moving on, therefore leaves the question of which composition differences can be trusted unanswered.

Here we present SLUICE (Screening Label Uncertainty to Identify Composition Effects), a workflow that treats each group difference as a claim to be tested for dependence on the labelling. SLUICE assigns each cell an identity using marker scores informed by a reference atlas, records the margin between the top-ranked and second-ranked identities, and applies three diagnostics: region-specific confidence thresholds, sensitivity analyses that reassign borderline calls, and single-gene driver checks (Figure 1). A difference is classed as robust if its direction is unchanged across the analyses and as fragile if it is not. SLUICE screens the annotation, not the Xenium measurement itself. We demonstrate the SLUICE workflow on Xenium Prime 5K data from mice with repetitive mild traumatic brain injury (rmTBI; hereafter TBI) or sham injury, in two regions. The optic tract was chosen because, in this injury model, it shows reduced axial diffusivity, altered functional connectivity, and increased Iba1 and GFAP immunoreactivity^7^, and the hippocampus because the effects of rmTBI there depend on age^8^ and for its relevance to memory and Alzheimer’s disease. The two regions differ in cellular makeup and therefore in how reliable the labels are, and we show which composition differences survive and which do not. We also discuss the current limitations of SLUICE, the next steps for validating it, and its possible future uses.

**Figure 1.**
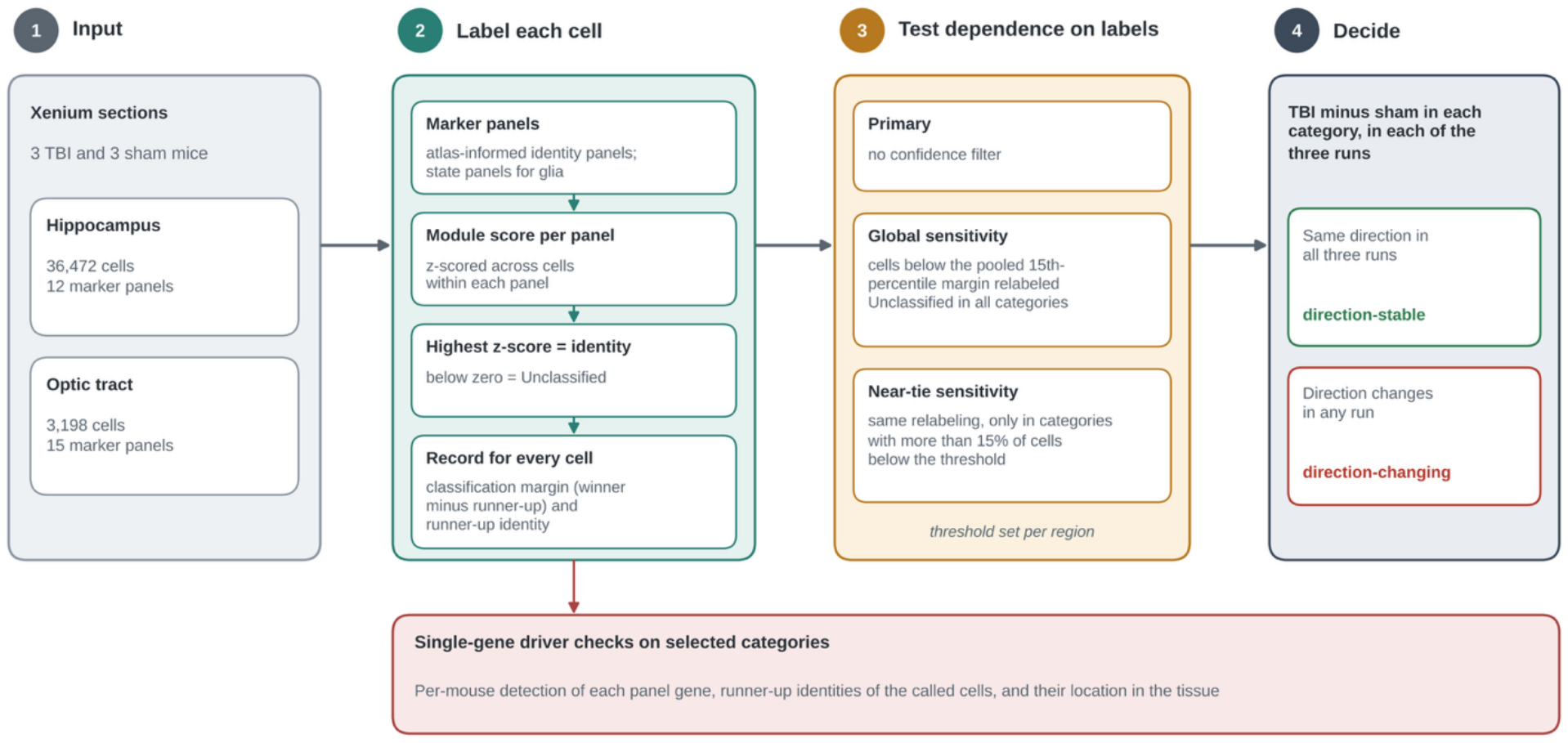
Overview of SLUICE. Cells are scored against marker panels and assigned the highest-scoring identity, and the classification margin and runner-up identity are recorded. Group differences are compared between the primary analysis and two sensitivity analyses, and single-gene driver checks are applied to selected categories.

## Methods

### Animals and repetitive mild traumatic brain injury

C57BL/6J mice were purchased from The Jackson Laboratory (Bar Harbor, ME). Animals were housed under standard conditions on a 12 h light/dark cycle with free access to food and water. All animal procedures were approved by the University of Florida Institutional Animal Care and Use Committee.

Female mice aged 2.5 months (n = 3 per group) were subjected to two mild closed-head injuries 24 h apart (0.6 J each) using the CHIMERA (Closed-Head Impact Model of Engineered Rotational Acceleration) device, or to a sham procedure. The injury and sham procedures, anesthesia, analgesia and euthanasia followed the protocol described previously^7^.

### Tissue preparation

At 7 days post-injury, mice were transcardially perfused with ice-cold sterile 0.9% saline and the brains were harvested. One hemisphere was fixed in 10% neutral buffered formalin at 4 °C for 24 h for spatial transcriptomics.

Formalin-fixed, paraffin-embedded (FFPE) coronal sections (5 µm thick) were mounted and prepared for the 10x Genomics Xenium In Situ platform according to the 10x Genomics Xenium FFPE Tissue Preparation Guide (CG000578).

### Xenium data acquisition

Sections from six mice (three TBI and three sham) were profiled by 10x Genomics through the Xenium Catalyst program using the Xenium Prime 5K Mouse Pan Tissue & Pathways panel (5,000 genes). Tissue sections from both the sham and rmTBI groups were mounted on a single specialized Xenium slide and processed according to the manufacturer’s (10x Genomics) instructions. Transcript detection, imaging, decoding, and onboard analysis were performed on the 10x Genomics Xenium Analyzer. Cells were segmented with the Xenium multimodal segmentation workflow: nuclei were identified from DAPI; cell boundaries from ATP1A1, E-cadherin, and CD45 staining; and interior stains (18S rRNA, αSMA, and vimentin) were used to refine the cell extent, with a nuclear expansion distance of 5 µm. The resulting datasets contained 46,403 to 72,905 cells per section, with a median of 478 to 574 transcripts and 354 to 408 genes per cell. The largest count includes a piece of cerebellum that was mounted with sections to identify the groups.

### Blinding

Samples were processed and analyzed using coded group labels to maintain blinding. Group identities were revealed after the blinded results had been presented to the laboratory on 30 September 2026. The marker panels, the 0.5% detection floor, the classification margin, and the Unclassified rule were established before unblinding. Four analysis decisions were made after unblinding: adoption of revised optic tract regions of interest (5 October 2026); the design of the confidence-filter sensitivity analyses (5 October 2026); exclusion of the choroid plexus panel from the hippocampus (6 October 2026); and exclusion of the neutrophil panel from the optic tract (8 October 2026). The two panel exclusions were based on the single-gene driver diagnostics described below, not on group differences; however, because they followed unblinding, they could be regarded as post hoc. The confidence threshold and near-tie categories were derived from margins pooled across all six mice without reference to group labels.

### Regions of interest

Regions of interest (ROIs) for the hippocampus and the optic tract were drawn as freehand polygons in Xenium Explorer (version 4.1.1) and exported as coordinate files, one per mouse and region. Cells were assigned to an ROI if their centroid fell inside the polygon (point-in-polygon test). To confirm that each polygon file was matched to the correct section, every run produced an overview figure showing each polygon drawn over the centroids of its sample (Figure 2A, B). Polygon areas were recorded, and cell density (cells per 1,000 µm^2^) was reported for each mouse, because ROI size and cell number differed between mice. Hippocampal ROIs were drawn by the first author before unblinding under the coded group labels. Optic tract ROIs were drawn by S.R., who was not blind to group, to include reactive and infiltrating cells at the border of the tract.

**Figure 2.**
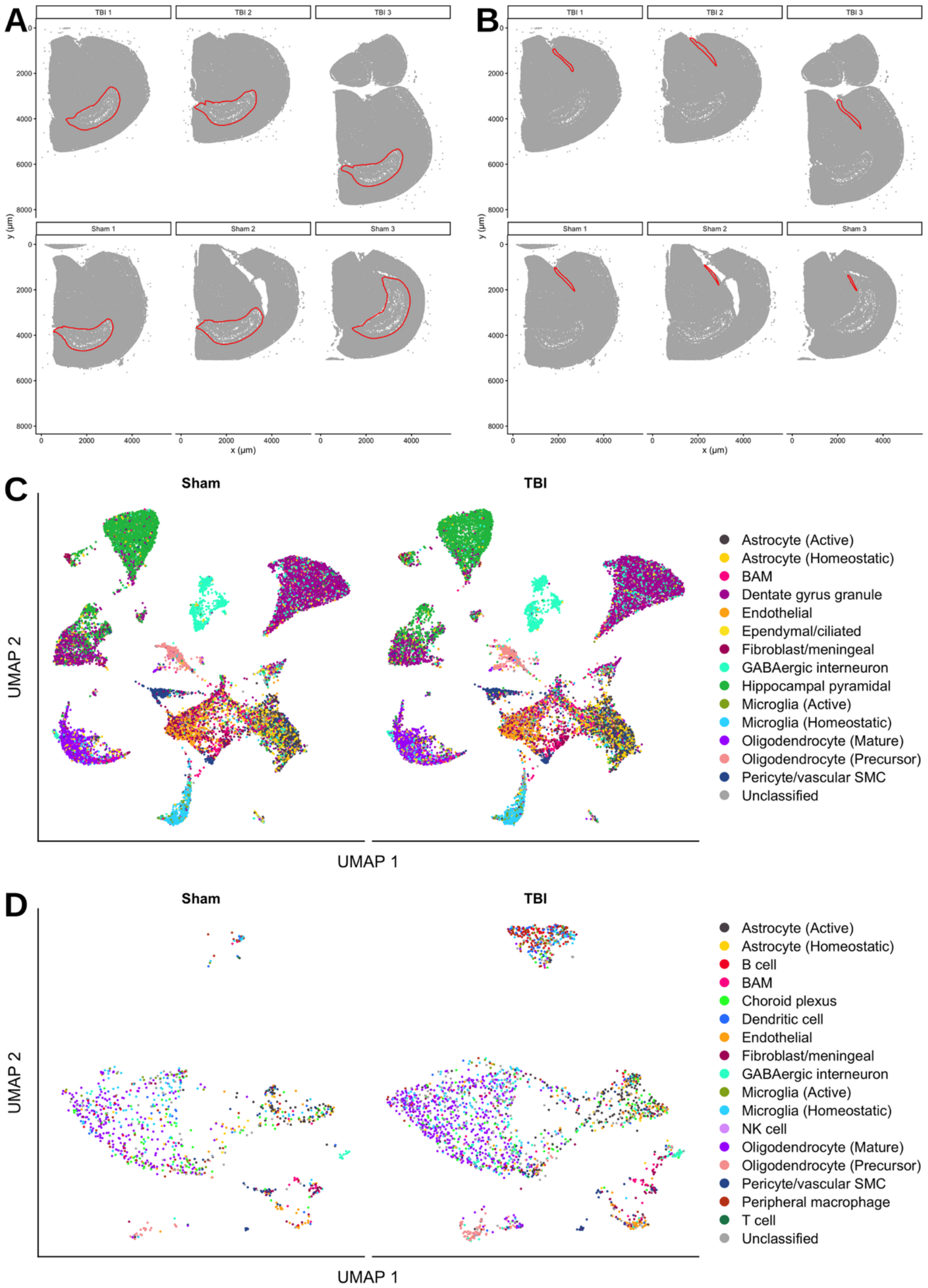
Regions of interest and cell-type labels. (A) Hippocampal regions of interest (red outlines) over all cells in each section. (B) Optic tract regions of interest, as in A. (C) UMAP of cell-type labels for the hippocampus from the primary analysis, with sham (19,380 cells) and TBI (17,092 cells) shown separately. (D) The same for the optic tract (sham, 960 cells; TBI, 2,238 cells). Cells are coloured by assigned identity. Unclassified cells are those for which no marker panel scored above the average (highest z-score below zero). UMAP axes are unitless embedding dimensions computed for display only. In A and B, y coordinates follow the Xenium Explorer orientation.

### Preprocessing and quality control

Xenium output was loaded into Seurat with LoadXenium function. Cells with fewer than 10 detected transcripts or fewer than 5 detected genes were removed, and counts were log-normalized (natural log) with Seurat’s NormalizeData function using default settings. As a check on the quality threshold, we recorded for each mouse the percentage of retained cells that would also pass a stricter cutoff of at least 20 transcripts and 10 genes; this exceeded 99.4% in every mouse in both regions.

### Cell-type classification

Cells were classified individually; no clustering was used to assign identities. Each cell was scored against a set of marker-gene panels, each containing one to eight genes. Identity panels were derived from the Allen Institute whole mouse brain taxonomy^9^ and checked gene by gene against the Xenium 5K panel; panels for border-associated macrophages and for cell states were curated from the literature^10–14^. The hippocampus was scored against 12 identity panels and the optic tract against 15 (see Region-specific panel sets).

Within each mouse, a panel was excluded from scoring if its most frequently detected gene was present in fewer than 0.5% of cells. For each remaining panel, a module score was computed with Seurat’s AddModuleScore function and z-scored across cells within that panel. Each cell was assigned the identity of the panel with the highest z-score; cells whose highest z-score was below zero were labeled Unclassified. For every cell, we recorded the classification margin, defined as the difference between the highest and second-highest panel z-scores, and the identity of the second-ranked (runner-up) panel.

Astrocytes, microglia, and oligodendrocyte-lineage cells were then assigned a state by rescoring them against two state panels and taking the higher module score (Table 1): active (Gfap, Serpina3n) versus homeostatic (Aldh1l1, Sox9) astrocytes; active (Trem2, Cst7, Lpl) versus homeostatic (Tmem119, Sall1, P2ry13) microglia; and mature (Mog, Mag, Cnp, Opalin) versus precursor (Pdgfra, Cspg4, Gpr17, Vcan) oligodendrocyte-lineage cells. A Purkinje cell panel (Pcp2) was included in both regions as a negative control and was expected to identify approximately 0% of cells; it fell below the detection floor in every mouse.

**Table 1.** Marker panels used to classify cells. Yes, the panel was scored in that region.

| Panel | Genes | Hippocampus | Optic tract |
|---|---|---|---|
| **Identity panels (first step)** |  |  |  |
| Astrocyte | Aldh1l1, Sox9 | Yes | Yes |
| Microglia | Tmem119, Sall1, P2ry13 | Yes | Yes |
| Oligodendrocyte lineage | Sox10, Olig1, Olig2 | Yes | Yes |
| Endothelial | Pecam1, Id1 | Yes | Yes |
| Pericyte/vascular SMC | Kcnj8, Myh11, Pdgfrb | Yes | Yes |
| Fibroblast/meningeal | Col1a1, Col1a2, Lum, Fn1 | Yes | Yes |
| BAM | Mrc1, F13a1, Cd163 | Yes | Yes |
| GABAergic interneuron | Gad1, Gad2, Slc32a1 | Yes | Yes |
| Hippocampal pyramidal | Slc17a7, Fibcd1, Spink8, Hes1, Rab38, Pou3f1, Hcn1, Map3k15 | Yes | – |
| Dentate gyrus granule | Slc17a7, Prox1 | Yes | – |
| Ependymal/ciliated | Foxj1, Pifo, Dnah12 | Yes | – |
| Choroid plexus | Folr1, Kl, Otx2 | – | Yes |
| Peripheral macrophage | Ccr2, Cd14, Ms4a7 | – | Yes |
| T cell | Cd3e, Cd3d, Cd3g | – | Yes |
| NK cell | Ncr1, Klrb1c, Gzmb, Klrk1, Prf1 | – | Yes |
| B cell | Cd19, Cd79a, Cd79b, Ms4a1, Cd22, Pax5 | – | Yes |
| Dendritic cell | Itgax, Cd83, Batf3, Xcr1, Clec9a | – | Yes |
| Purkinje (negative control) | Pcp2 | Yes | Yes |
| **State panels (second step)** |  |  |  |
| Astrocyte, active | Gfap, Serpina3n | Yes | Yes |
| Astrocyte, homeostatic | Aldh1l1, Sox9 | Yes | Yes |
| Microglia, active | Trem2, Cst7, Lpl | Yes | Yes |
| Microglia, homeostatic | Tmem119, Sall1, P2ry13 | Yes | Yes |
| Oligodendrocyte, mature | Mog, Mag, Cnp, Opalin | Yes | Yes |
| Oligodendrocyte, precursor | Pdgfra, Cspg4, Gpr17, Vcan | Yes | Yes |

Identity panels were scored for every cell. State panels were applied in a second step to cells already assigned to the astrocyte, microglia, or oligodendrocyte-lineage identity. Purkinje (Pcp2) is a negative control. The choroid plexus panel was evaluated in the hippocampus and excluded after unblinding. Neutrophil (Ly6g, Csf3r, Cxcr2) and retinal ganglion cell (Pou4f1, Sncg) panels were evaluated in the optic tract and excluded (see Results and Methods). Identity panels were derived from the Allen Institute whole-mouse-brain taxonomy<sup>9</sup> and checked gene by gene against the Xenium 5K panel; the BAM and state panels were curated from the literature, and the five optic tract immune panels were assembled by the first author from canonical markers of each cell type and checked for detection in the Xenium panel.

### Region-specific panel sets

Panels were included or excluded per region on the basis of anatomy and the driver-gene diagnostics described below. The optic tract was scored against 15 panels: the 13 original panels minus hippocampal pyramidal, dentate gyrus granule and ependymal/ciliated, plus five immune panels (peripheral macrophage, T cell, NK cell, B cell and dendritic cell). Neutrophil and retinal ganglion cell panels were also evaluated for the optic tract and excluded; the evidence for each exclusion is reported in Results. The hippocampus was scored against the 13 original panels minus choroid plexus, giving 12.

### Classification confidence and sensitivity analysis

The primary analysis applied no confidence filter. To test whether group differences depended on borderline calls, we ran two sensitivity analyses per region. For each region, the 15th percentile of the classification margin, pooled across all six mice, defined a confidence threshold (hippocampus, 0.17; optic tract, 0.026). In the first sensitivity analysis (global), cells with margins below this threshold were relabeled Unclassified in every category. In the second (near-tie), relabeling was restricted to categories in which more than 15% of cells in the primary run fell below the threshold (hippocampus: border-associated macrophages, ependymal/ciliated cells and active microglia; optic tract: choroid plexus, GABAergic interneurons, pericytes/vascular smooth muscle cells, B cells and peripheral macrophages). The threshold and the near-tie categories were derived without reference to group labels. The 15th percentile was chosen during development on the optic tract and applied unchanged to both regions.

A group difference in composition was considered robust to near-tie handling if the direction of the TBI-minus-sham difference was the same in all three runs (primary, global and near-tie); the size of each shift was also reported.

### Single-gene driver diagnostics

Confidence margins cannot detect a call that is confidently wrong. We therefore examined categories that were biologically implausible in a region, or that were unstable in the sensitivity analyses, with three complementary checks. First, gene co-detection: for each mouse, the percentage of cells assigned to the category in which each of its panel genes was detected, compared with all other cells. A genuine cell type is expected to express its panel genes together; detection of one gene in most calls with the others near zero indicates a single-gene driver. Second, runner-up identity: the distribution of runner-up panels among the category’s cells, pooled across mice, together with their median classification margin. Third, spatial location: the position of the category’s cells within the ROI, plotted against a reference cell type (pyramidal and dentate gyrus granule neurons in the hippocampus; mature oligodendrocytes in the optic tract). Categories whose calls were driven by a single gene, had runner-up identities indicating another cell type, and were located inconsistently with their expected anatomy were excluded from that region’s panel set, and the region was reanalyzed. Earlier runs were retained for comparison.

### Composition statistics

For each mouse, the percentage of cells in each category was calculated, with categories absent from a mouse recorded as 0%. Group means were compared with Welch’s t-test (chosen because equality of variances cannot be assessed with three animals per group), with the mouse as the unit of analysis and Benjamini-Hochberg correction across all categories tested in a region. With three animals per group, these comparisons were treated as exploratory.

### Visualization

For display only, the samples in each region were merged, and principal components (30) and a UMAP embedding were computed with Seurat (Figure 2C, D); these were not used to assign identities.

### Software, data and code availability

Analyses were performed in R (version 4.6.1) with Seurat (version 5.5.1)^5^, dplyr (version 1.2.1) and ggplot2 (version 4.0.3); a full list of package versions is provided in the repository. All analysis code and the summary tables underlying the figures are available at https://github.com/rebeccasakko/hippocampus-xenium-analysis. The raw Xenium data are part of ongoing studies and are available from the corresponding author upon reasonable request.

## Results

### Region-specific confidence thresholds

In each region, we set the confidence threshold at the 15th percentile of the classification margin across all cells, where the margin is the score of the winning label minus that of the runner-up. The threshold was 0.17 in the hippocampus (12 marker panels, 36,472 cells) and 0.026 in the optic tract (15 panels, 3,198 cells), a difference of almost sevenfold (Figure 3). A cutoff chosen in one region would therefore relabel a very different share of cells in the other. Applied to the optic tract, the hippocampal cutoff would have relabeled 27.1% of cells as Unclassified instead of 15%, whereas applied to the hippocampus the optic tract cutoff would have relabeled 4.7%.

**Figure 3.**
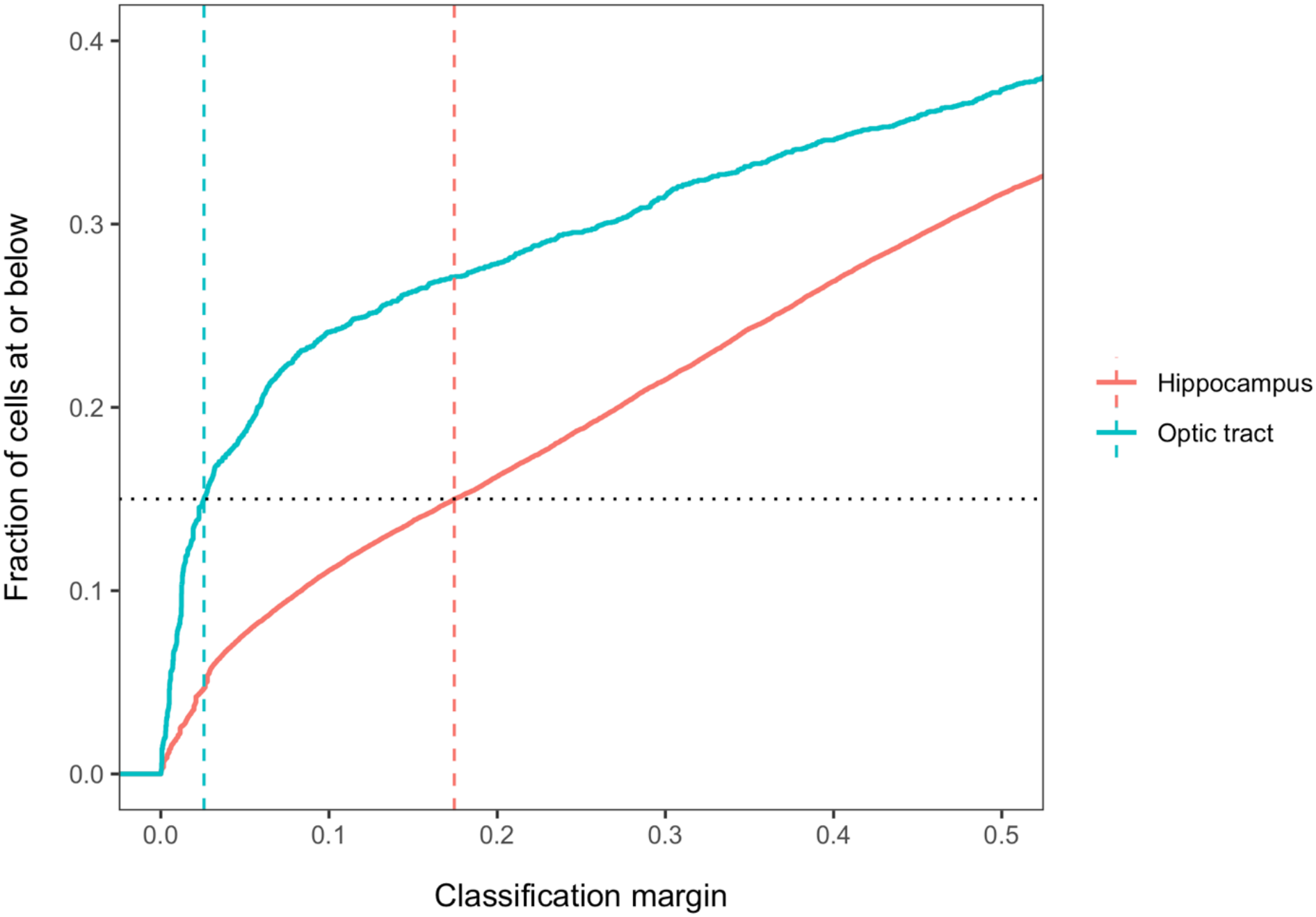
Region-specific confidence thresholds. Cumulative distribution of the classification margin in the hippocampus (36,472 cells) and the optic tract (3,198 cells). Dashed lines mark each region’s 15th-percentile threshold (0.17 and 0.026); the dotted line marks the 15% level.

### Single-gene driver checks

A margin describes how close a call was, not whether the winning label is correct. For selected categories, we therefore checked which panel genes were detected in the called cells, which label was the runner-up, and where the cells lay in the tissue (Figure 4; Tables 2 and 3).

**Table 2.** Detection of choroid plexus panel genes in hippocampal cells called choroid plexus.

| **Mouse** | **Cells called** | **Folr1** | **Kl** | **Otx2** |
|---|---|---|---|---|
| TBI 1 | 245 | 1.2 | 82.9 | 1.6 |
| TBI 2 | 306 | 2.0 | 95.8 | 1.0 |
| TBI 3 | 397 | 1.8 | 69.3 | 1.3 |
| Sham 1 | 370 | 1.6 | 65.7 | 1.4 |
| Sham 2 | 351 | 0.6 | 90.6 | 2.0 |
| Sham 3 | 428 | 0.7 | 98.4 | 0.9 |
| *Outside the category (range)* |  | *0.0 to 0.1* | *0.2 to 0.3* | *0.0 to 0.1* |

Values are the percentage of called cells in which each gene was detected, from the run that included the choroid plexus panel. The last row gives the range across mice of the percentage of all other cells in which the gene was detected. Mice are labeled by group (TBI or sham) and number.

**Table 3.** Detection of neutrophil panel genes in optic tract cells called neutrophils.

| **Mouse** | **Cells called** | **Ly6g** | **Csf3r** | **Cxcr2** |
|---|---|---|---|---|
| TBI 1 | 24 | 0.0 | 100.0 | 4.2 |
| TBI 2 | 29 | 0.0 | 96.6 | 0.0 |
| TBI 3 | 38 | 0.0 | 100.0 | 0.0 |
| Sham 1 | 79 | 0.0 | 5.1 | 0.0 |
| Sham 2 | 4 | 0.0 | 100.0 | 0.0 |
| Sham 3 | 2 | 0.0 | 100.0 | 0.0 |
| *Outside the category (range)* |  | *0.0* | *0.3 to 7.9* | *0.0* |

Values are the percentage of called cells in which each gene was detected, from the run that included the neutrophil panel. The last row gives the range across mice of the percentage of all other cells in which the gene was detected. Sham 2 and Sham 3 had 4 and 2 called cells, so their percentages rest on very few cells.

**Figure 4.**
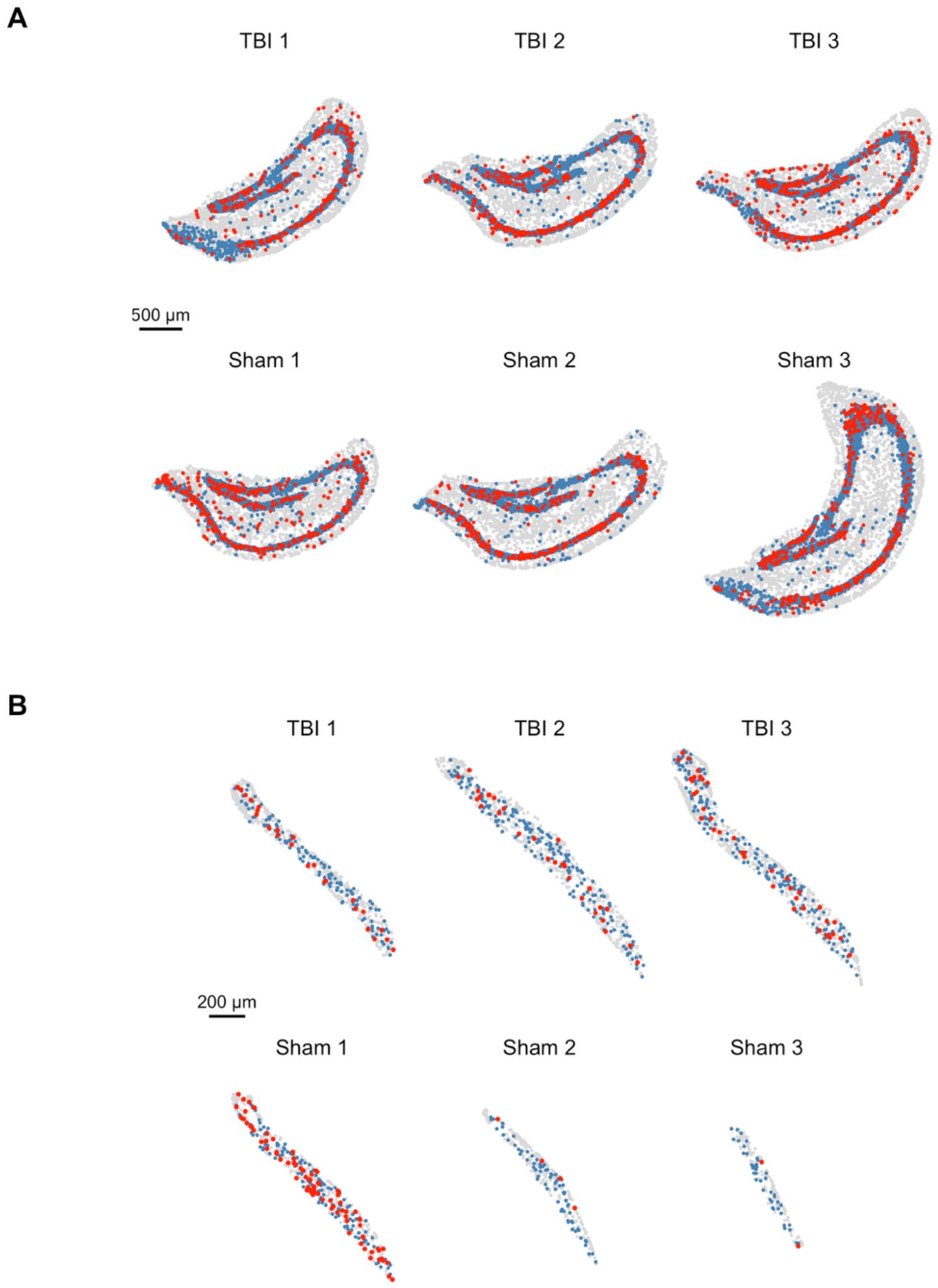
Single-gene driver checks. (A) Location of cells called choroid plexus in the hippocampus (red) against hippocampal pyramidal and dentate gyrus granule cells (blue). (B) Location of cells called neutrophils in the optic tract (red) against mature oligodendrocytes (blue). Grey points are all other cells. Panels are drawn at the same scale within each region; scale bars, 500 µm (A) and 200 µm (B). Detection of the panel genes in these cells, per mouse, is given in Tables 2 and 3.

#### Hippocampus, choroid plexus

In the run that included the choroid plexus panel, 2,097 hippocampal cells (5.7%) were called choroid plexus, 245 to 428 per mouse. Kl (Klotho) was detected in 66% to 98% of these cells in every mouse (Table 2), whereas the other two panel genes, Folr1 and Otx2, were each detected in 2.0% or fewer. Outside the category, Kl was detected in about 0.2% of cells. For 74% of the called cells, the runner-up was a neuronal label (46% hippocampal pyramidal, 24% dentate gyrus granule, and 5% GABAergic interneuron), and the cells lay along the pyramidal and dentate cell layers. The category was therefore collecting Kl-expressing cells that were not choroid plexus, and the panel was excluded from the hippocampus after unblinding (6 October 2026). Relabelling low-margin cells as Unclassified changed the choroid plexus group difference by only 0.12 percentage points across the primary, global and near-tie runs (−0.40, −0.52 and −0.40), so margin-based filtering left these calls largely in place.

#### Optic tract, neutrophils

In the optic tract, 176 cells were called neutrophils, from 2 to 79 per mouse. In five of six mice, Csf3r was detected in 97% to 100% of the called cells (Table 3), but Ly6g was detected in none and Cxcr2 in no more than 4.2% in any mouse. The exception was one sham mouse with 79 called cells, of which only 5% carried Csf3r. Runner-up labels were largely other myeloid types (peripheral macrophage 18%, dendritic cell 13%, and microglia 13%), together with choroid plexus (32%). The median margin of the neutrophil calls was 0.21, about eight times the optic tract threshold of 0.026, and the cells were scattered evenly through the tract, so margin filtering would not have removed them. These cells are best described as Csf3r-expressing myeloid cells and not neutrophils, and the neutrophil panel was excluded from the optic tract after unblinding (8 October 2026).

#### Robust and fragile composition differences

We applied the screen to the composition results in each region. For every category, we calculated the TBI-minus-sham difference in the mean percentage of cells under the primary analysis and under the global and near-tie sensitivity analyses (Figure 5; Tables 4 and 5). A difference was classed as direction-stable if its sign was the same in all three runs, and as direction-changing otherwise; the largest shift in its size between runs is also shown.

**Table 4.** Hippocampus: TBI-minus-sham difference in composition under the primary and two sensitivity analyses.

| **Category** | **Primary** | **Global** | **Near-tie** | **Largest shift** | **Direction** |
|---|---|---|---|---|---|
| Ependymal/ciliated | −1.01 | +0.38 | +0.38 | 1.39 | **Changes** |
| BAM | +1.29 | +0.08 | +0.08 | 1.21 | Stable |
| Unclassified | −0.07 | −0.86 | −0.15 | 0.79 | Stable |
| Fibroblast/meningeal | −0.35 | +0.12 | −0.35 | 0.47 | **Changes** |
| Dentate gyrus granule | −0.79 | −0.38 | −0.79 | 0.41 | Stable |
| Hippocampal pyramidal | −1.17 | −0.85 | −1.17 | 0.32 | Stable |
| Pericyte/vascular SMC | +0.30 | −0.01 | +0.30 | 0.31 | **Changes** |
| GABAergic interneuron | −0.29 | −0.44 | −0.29 | 0.15 | Stable |
| Astrocyte (Homeostatic) | +0.84 | +0.70 | +0.84 | 0.14 | Stable |
| Endothelial | +0.46 | +0.56 | +0.46 | 0.10 | Stable |
| Microglia (Active) | +0.22 | +0.12 | +0.12 | 0.10 | Stable |
| Astrocyte (Active) | −0.06 | −0.14 | −0.06 | 0.08 | Stable |
| Oligodendrocyte (Precursor) | −0.23 | −0.16 | −0.23 | 0.07 | Stable |
| Oligodendrocyte (Mature) | +0.72 | +0.76 | +0.72 | 0.04 | Stable |
| Microglia (Homeostatic) | +0.14 | +0.11 | +0.14 | 0.03 | Stable |

Values are TBI minus sham in mean percentage of cells (percentage points); positive values are higher in TBI. Primary, no confidence filter; Global, cells below the pooled 15th-percentile margin relabeled Unclassified in all categories; Near-tie, the same relabeling restricted to categories with more than 15% of cells below the threshold. The largest shift is the range of the three values. Direction is Stable if the sign is the same in all three runs. n = 3 mice per group.

**Table 5.** Optic tract: TBI-minus-sham difference in composition under the primary and two sensitivity analyses.

| Category | Primary | Global | Near-tie | Largest shift | Direction |
|---|---|---|---|---|---|
| Unclassified | +4.31 | −1.17 | −0.12 | 5.48 | **Changes** |
| B cell | −0.19 | +2.85 | +2.85 | 3.04 | **Changes** |
| Peripheral macrophage | +0.60 | +3.05 | +3.05 | 2.45 | Stable |
| GABAergic interneuron | +1.76 | +0.83 | +0.83 | 0.93 | Stable |
| Pericyte/vascular SMC | −0.16 | −0.95 | −0.95 | 0.79 | Stable |
| Dendritic cell | +1.03 | +1.75 | +1.03 | 0.72 | Stable |
| Choroid plexus | −1.19 | −0.53 | −0.53 | 0.66 | Stable |
| Microglia (Active) | −0.67 | −0.29 | −0.67 | 0.38 | Stable |
| BAM | −0.18 | −0.44 | −0.18 | 0.26 | Stable |
| Fibroblast/meningeal | −0.37 | −0.16 | −0.37 | 0.21 | Stable |
| Astrocyte (Active) | +0.55 | +0.35 | +0.55 | 0.20 | Stable |
| Microglia (Homeostatic) | −1.71 | −1.51 | −1.71 | 0.20 | Stable |
| Astrocyte (Homeostatic) | −1.15 | −1.26 | −1.15 | 0.11 | Stable |
| Endothelial | −1.24 | −1.17 | −1.24 | 0.07 | Stable |
| Oligodendrocyte (Mature) | −2.14 | −2.11 | −2.14 | 0.03 | Stable |
| Oligodendrocyte (Precursor) | +1.27 | +1.30 | +1.27 | 0.03 | Stable |
| NK cell | −0.21 | −0.21 | −0.21 | 0.00 | Stable |
| T cell | −0.32 | −0.32 | −0.32 | 0.00 | Stable |

Values are TBI minus sham in mean percentage of cells (percentage points); positive values are higher in TBI. Primary, no confidence filter; Global, cells below the pooled 15th-percentile margin relabeled Unclassified in all categories; Near-tie, the same relabeling restricted to categories with more than 15% of cells below the threshold. The largest shift is the range of the three values. Direction is Stable if the sign is the same in all three runs. n = 3 mice per group.

**Figure 5.**
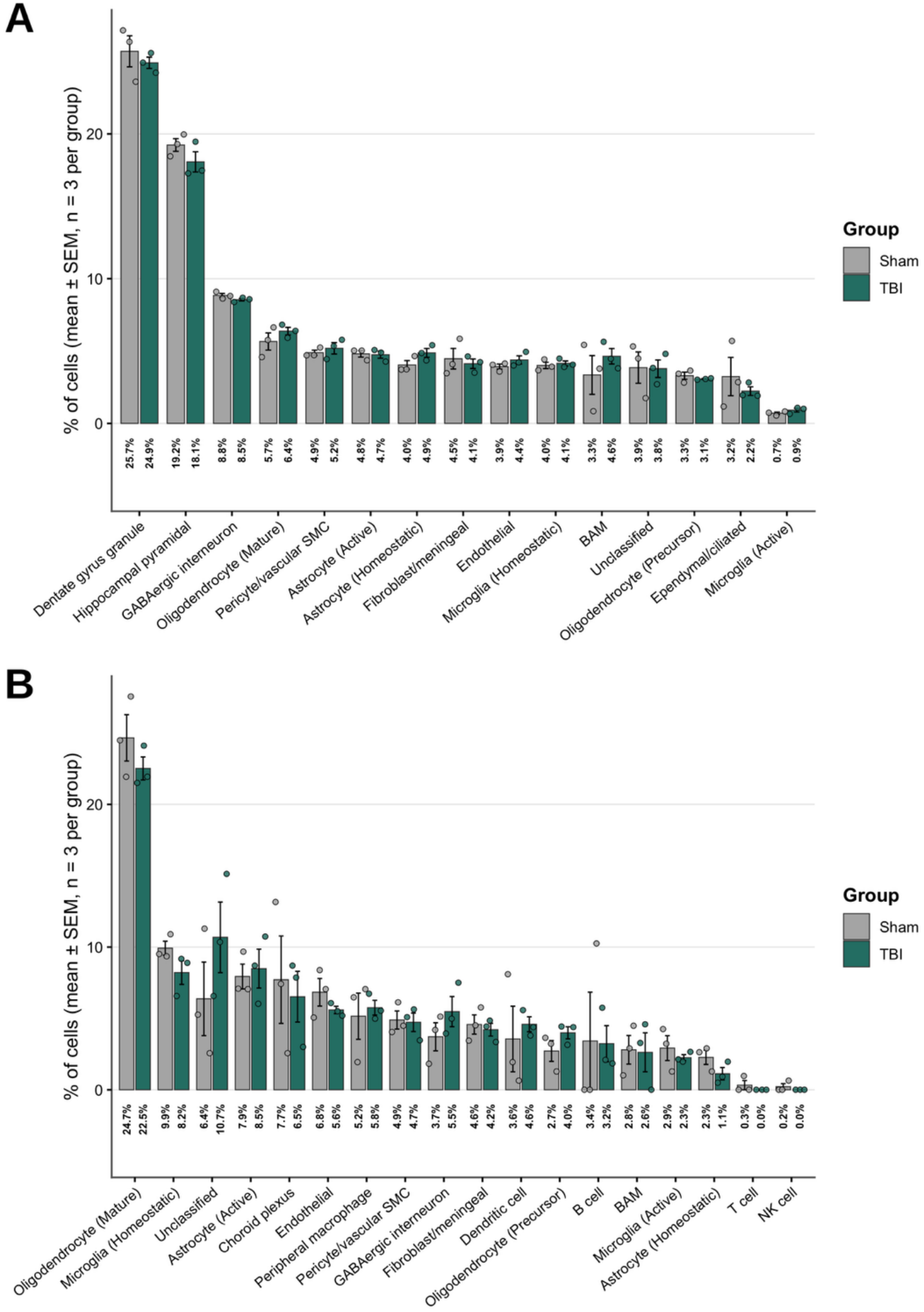
Cell-type composition in the primary analysis. Percentage of cells assigned to each category in sham (grey) and TBI (teal) mice for the hippocampus (A) and the optic tract (B). Bars show the group means, and error bars show the SEM (n = 3 mice per group); circles show individual mice. Values under the bars are group means. Categories are ordered by overall mean abundance. Differences under the sensitivity analyses are given in Tables 4 and 5.

In the hippocampus (12 panels; confidence threshold 0.17), 12 of 15 categories were direction-stable. Hippocampal pyramidal and dentate gyrus granule cells were lower, and homeostatic astrocytes and mature oligodendrocytes were higher, in TBI in every run. Three categories changed direction: ependymal/ciliated cells (−1.01 to +0.38 percentage points), and fibroblast/meningeal and pericyte/vascular smooth muscle cells, the last two only in the global run and from differences close to zero. BAM kept its direction, but its difference fell from +1.29 to +0.08 percentage points once low-margin calls were relabeled. In five of the six mice, only 16–31% of cells called BAM in the primary run carried a transcript of any BAM marker gene (Mrc1, F13a1, or Cd163), so the primary BAM difference reflected the number of marker-free calls and not BAM abundance.

In the optic tract (15 panels; threshold 0.026), 16 of 18 categories were direction-stable. B cells (−0.19 to +2.85) and Unclassified (+4.31 to −1.17 and −0.12) changed direction. Peripheral macrophages kept their direction, but the difference grew from +0.60 to +3.05 percentage points, and the choroid plexus difference roughly halved (−1.19 to −0.53). The main glial and vascular populations (mature oligodendrocytes, homeostatic microglia, homeostatic astrocytes, precursor oligodendrocytes and endothelial cells) changed by no more than 0.2 percentage points across runs.

No category differed between groups after Benjamini–Hochberg correction in either region (smallest adjusted P, 0.74 in the hippocampus and 0.84 in the optic tract; n = 3 per group). Robustness to near-tie handling is therefore a necessary condition for carrying a difference forward, not evidence that the difference is significant.

## Discussion

SLUICE addresses a practical problem in small spatial transcriptomics studies: a difference in cell-type composition can depend on how borderline cells were labelled. In both regions, most categories kept their direction across the primary and two sensitivity analyses, and a minority did not, so the screen sorts differences into those that can be carried forward and those that need caution. The confidence threshold differed almost sevenfold between the hippocampus (0.17) and the optic tract (0.026), which shows that a cutoff chosen in one tissue should not be applied to another. No category differed between groups after correction for multiple testing in either region (n = 3 per group). SLUICE is therefore a way to prioritise differences for follow-up, and it does not test them. A difference that survives the screen is not shown to be real, and one that fails it should not be reported without that caveat.

The driver checks found errors that confidence margins could not. Neurons expressing Kl were called choroid plexus in the hippocampus, and myeloid cells expressing Csf3r were called neutrophils in the optic tract. For the choroid plexus calls, margin-based filtering changed the group difference by only 0.12 percentage points, so it did not remove the error, and the neutrophil calls had a median margin about eight times the optic tract threshold. Excluding the neutrophil panel from the optic tract reduced the number of categories whose difference changed direction from five to two. The exclusion of the choroid plexus panel (hippocampus) and of the neutrophil panel (optic tract) were decisions made after unblinding and are reported as such in the Methods; they were made based on the gene-level evidence and not on the group difference.

The BAM category shows why the size of a difference matters as well as its direction. In the hippocampus, the BAM difference kept its sign but fell from +1.29 to +0.08 percentage points once low-margin calls were relabelled. Its marker genes were detected in fewer than 1% of cells, yet about 4–5% of cells were called BAM, and in five of six mice only 16–31% of BAM calls carried a transcript of any BAM marker. Atlas-informed scoring will always return a winner, and a category with few detectable markers can collect cells that fit no other label well. A direction-only rule would have classed BAM as stable, which is why the tables also report the largest shift between runs. We did not set a size cutoff for calling a shift too large, because with three animals per group we had no basis for choosing one.

The study has several limitations. The comparison rests on three mice per group, so the statistical tests have little power, and the absence of a significant difference should not be read as the absence of an effect. The data come from one platform and one gene panel, and the classification depends on marker lists, which were drawn from atlases and from knowledge of the tissue and could be chosen differently. The sensitivity analyses relabel cells as Unclassified, and Unclassified itself changed direction in the optic tract, so that category must be read with the others. Several analysis decisions were made after unblinding, although the marker panels, detection floor, and margin rule were fixed before it. The optic tract regions of interest were drawn by a co-author who knew the group identities, whereas the hippocampal regions were drawn blind, and the sections may not have been cut at identical anatomical levels; one sham hippocampal section was noticeably larger than the others, and differences in level can shift composition independently of injury. Composition percentages within a mouse sum to 100% and are not independent, but we compared categories one at a time; methods designed for compositional data, such as propeller^15^, would be preferable in larger studies. The five optic tract immune panels were assembled from canonical markers and not taken from a published atlas, and the 0.5% detection floor was chosen pragmatically rather than optimized. We did not validate any label by an independent method such as immunostaining, so the driver checks identify likely errors but do not prove them. SLUICE also screens the annotation and cannot detect problems in the measurement or segmentation themselves.

The most direct next step is validation at the bench. In situ hybridization or immunostaining for the genes behind the fragile and suspected mislabelled calls, such as Kl in the hippocampal cells called choroid plexus and Csf3r and Ly6g in the optic tract cells called neutrophils, would test whether these cells are what the labels say, and we plan this validation in follow-up work. Future work should also test SLUICE on public spatial datasets with larger group sizes, compare its calls with those of reference-mapping methods, and extend the comparison step to mixed-effects models or pseudobulk differential expression that make fuller use of the animal-level structure^16,17^. For now, we suggest that studies of this kind report, for each cell type, whether its group difference is stable across reasonable labelling choices, and set confidence thresholds for each region rather than for the whole experiment.

## Data and code availability

The raw Xenium data are part of ongoing studies and are not publicly available at this time; they are available from the corresponding author upon reasonable request. All analysis code and the summary tables underlying the figures are available at https://github.com/rebeccasakko/hippocampus-xenium-analysis.

## Author contributions

R.S.: conceptualization, methodology, software, formal analysis, writing (original draft). S.R.: investigation (rmTBI surgeries, tissue sectioning), resources, supervision. H.E.H.: software, validation. J.F.A.: conceptualization, supervision, funding acquisition, writing (review and editing).

## Acknowledgements

This work was supported by the National Institute on Aging of the National Institutes of Health under award numbers R01AG074584, R01AG075900 and R21AG095654. The content is solely the responsibility of the authors and does not necessarily represent the official views of the National Institutes of Health. We thank 10x Genomics for providing the Xenium Prime 5K run through the Xenium Catalyst program.

Use of AI tools. The AI assistant Claude (Anthropic) was used to help write and debug the analysis code, to help assemble the candidate marker genes for the optic tract immune panels, and to help draft and edit the manuscript text. The authors reviewed and verified all code, marker panels, and text, and take full responsibility for the content of this article.

